# Explicitly Nonlinear Dynamic Functional Network Connectivity In Resting-State fMRI Data

**DOI:** 10.1101/2022.06.22.497262

**Authors:** S. M. Motlaghian, A. Belger, J. R. Bustillo, A. Faghiri, J. M. Ford, A. Iraji, K. Lim, D. H. Mathalon, R. Miller, B. A. Mueller, D. O’Leary, G. Pearlson, S. G. Potkin, A. Preda, T.G. van Erp, V. D. Calhoun

## Abstract

Most dynamic functional connectivity in fMRI data is focused on linear correlations, and to our knowledge, no study has studied whole brain explicitly nonlinear dynamic relationships within the data. While some approaches have attempted to study overall connectivity more generally using flexible models, we are particularly interested in whether the non-linear relationships, above and beyond linear, are capturing unique information. This study thus proposes an approach to assess the explicitly nonlinear dynamic functional network connectivity derived from the relationship among independent component analysis time courses. Linear relationships were removed at each time point to evaluate, typically ignored, explicitly nonlinear dFNC using normalized mutual information. Simulations showed the proposed method accurately estimated NMI over time, even within relatively short windows of data. Results on fMRI data included 151 schizophrenia patients, and 163 healthy controls showed three unique, highly structured, mostly long-range, functional states that also showed significant group differences. This analysis identifies a higher level of explicitly nonlinear dependencies in transient connectivity within the visual network in healthy controls compared to schizophrenia patients. In particular, nonlinear relationships tend to be more widespread than linear ones. We also find highly significant differences in the relative co-occurrence of linear and explicitly nonlinear states in HC and SZ, suggesting these may be an important aspect of the disorder. Overall, this work suggests that quantifying nonlinear dependencies of dynamic functional connectivity may provide a complementary and potentially valuable tool for studying brain function by exposing relevant variation that is typically ignored.

## Introduction

Functional connectivity (FC) and its network analog, functional network connectivity (FNC), are widely used to study whole brain resting brain function. These methods study the relationship between time courses (TC) from different brain regions or networks (E. Allen et al., 2011; Bastos & Schoffelen, 2016; Friston, 2011; Sala-Llonch, Bartrés-Faz, & Junqué, 2015; van den Heuvel & Hulshoff Pol, 2010). Other research has focused on modeling brain activity using nonlinear models (Lahaye, Poline, Flandin, Dodel, & Garnero, 2003; Stam, 2005; Su, Wang, Shen, Feng, & Hu, 2013; Wismüller, Wang, Dsouza, & Nagarajan, 2014). The nonlinear effects of hemodynamic responses in fMRI data (Deneux & Faugeras, 2006; Miller et al., 2001; Obata et al., 2004), which, crucially, can also vary with time (and location) and changes from subject to subject (de Zwart et al., 2009). Considering even just these few examples of nonlinear effects, it is likely, even expected, that distinct brain areas might be nonlinearly related in a way that would be missed by conventional linear F(N)C analysis. Our prior work in this direction indicated a modular nonlinear FNC between whole brain networks (Motlaghian et al., 2021).

The work mentioned thus far is all focused on static functional connectivity measuring temporal coherence averaged across the entire experiment. More recent studies have focused on assessing the dynamics in FNC (dFNC) over time to capture additional insight into the underlying properties of brain activities. One way to approach this is to divide the time courses into smaller windows and measure the temporal coherence between signals within each successive window. This method is known as the sliding window approach (E. A. Allen et al., 2014; Hindriks et al., 2016; Lindquist, Xu, Nebel, & Caffo, 2014) and is widely used in the field. However, virtually all time-resolved whole brain approaches have focused only on time-varying linear dependencies among networks or regions.

The current work is motivated by our prior work (Motlaghian et al., 2021) which identified informative and highly structured explicitly nonlinear relationships in static FNC. Here, our focus is to extend our previous approach to study the dynamic of explicitly nonlinear (EN) dependencies between ICN’s time courses and evaluate the properties of these relationships. That is, we are interested in studying the nonlinear information above and beyond the linear effects, i.e., what is typically ignored in a linear analysis. We first propose an approach to capture explicitly nonlinear information by removing the linear relationships identified within a sliding window approach, then analyzing the residual, explicitly nonlinear relationships using normalized mutual information (NMI). We first show our approach works well within simulated data, even though we are estimating NMI over relatively short windows of time. For real fMRI data, we extract the transient nonlinear patterns of FNC and then we evaluate whether these activities are different in controls (HC) and patients with schizophrenia (SZ). We present the approach in section 2.3, then demonstrate it via simulation in section 2.4. Section 2.5 explained how the proposed method is applied to resting-state fMRI data, including 163 controls and 151 patients. We also compute the linear dFNC and compare the result with nonlinear dFNC findings in section 2.6.

Results showed three distinctive and highly structured EN dynamic states. Several evaluations, such as fractional occupancy, dwell time, and state transition probability, are performed to study how brain contributes to each state (Section 3.2). The interpretation of these analyses indicates a high level of linear and nonlinear dependency coefficients within and between networks in controls compared to individuals with schizophrenia. We also find significant differences in the concurrence of the linear and explicitly nonlinear states, again highlighting the importance of capturing such, typically ignored, information.

## Materials and Methods

### 2.1. Quantifying Explicitly Nonlinear Dependency via a Normalized Mutual Information Approach

The main aim of this work is to estimate the dynamic of EN dependencies among ICNs, using a sliding window analysis approach (E. A. Allen et al., 2014; Hutchison, Womelsdorf, Allen, et al., 2013; Hutchison, Womelsdorf, Gati, Everling, & Menon, 2013; Saha et al., 2020). For each pair x and y of ICNs, we first estimate the linear correlation measured by a linear model 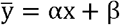, where 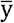 is the best linear fit predicting y given x, α is the slope and β is the vertical intercept. Next, the linear effect is removed by calculating 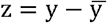. Then dependencies between x and z is measured by NMI. The formula for calculating the value of NMI is

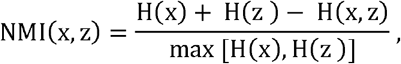

where H(x) and, H(z) are marginal entropies and H(x, z) is the joint entropy. The NMI measurement can values between 0 and 1, if it has a value of 0 this means there is no dependency between x and z, and 1 indicates an absolute dependence of two variables.

We apply the same method for assessing the dynamics of the nonlinear dependencies. Let x_t_ and y_t_ represent samples of x and y in the window t. Then by linear regression 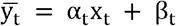, we estimate linear relation between x_t_ and y_t_. Next, the linear effect is removed by calculating 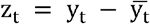. Lastly, dependencies between x_t_ and z_t_ is measured by NMI. Swapping x_t_ and y_t_ may result in a slightly different value. Thus, we consider the average of both results to ensure symmetry.

### 2.2. Simulated Experiment

The length of the sliding window (number of time points in the window) needed to be wide enough to ensure a valuable estimation of nonlinear dependencies. This size is selected by measuring dependencies in simulated data by NMI. The decisive length is determined by the criteria where NMI can successfully distinguish nonlinear dependency from linear dependency. The impact of the shape of the relationship and the number of sample points on NMI are studied in this simulation.

Our focus for the shape was on linear and nonlinear dependencies. To do so, we modeled three types of relationships. We created a vector x of size 1000 × 1 where its components are generated from a random uniform distribution on [0 1], for three cases as follows:

I. Vector y_1_ has a purely linear relationship with x.
II. Vector y_2_ has a quadratic relationship and no linear correlation with x. That is x and y_2_ have only a nonlinear dependency.
III. Vector y_3_ has a combination of linear and nonlinear correlation with x.

Gaussian noise of zero mean is added to each equation and plotted in Figure. 1Figure. 1.

**Figure. 1:**
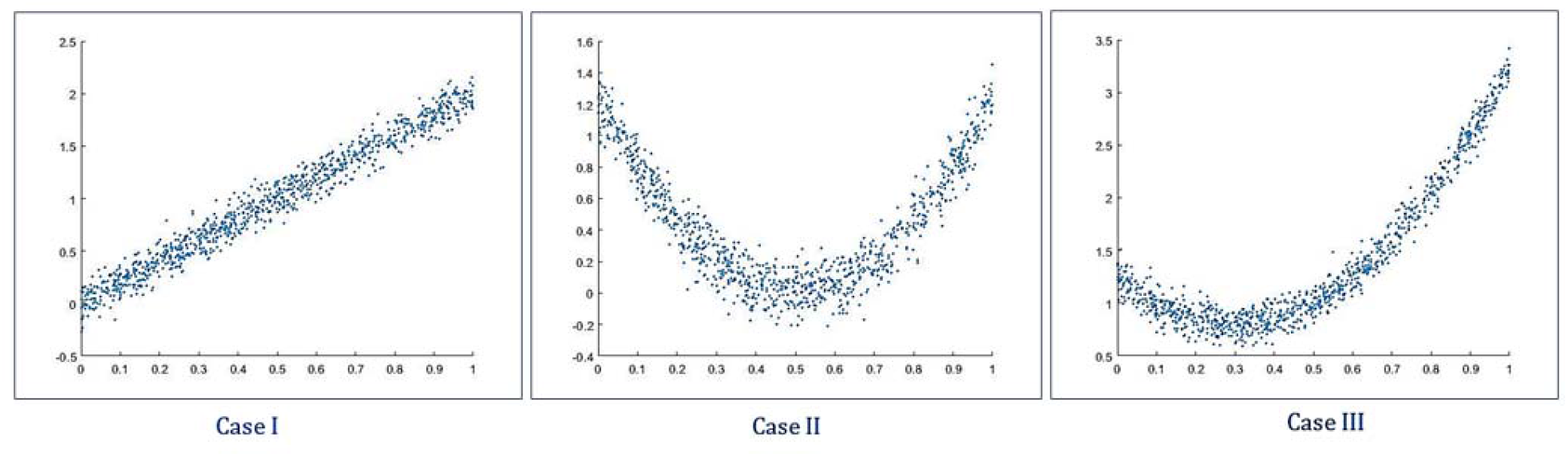
Three simulation cases for linear and nonlinear correlation between two vectors. Vector *x* has its components randomly derived from a uniform distribution [0 1]. From left to right, we have Case I, Case II, and Case III such that in Case I, *y*_l_ = 2*x* + *ε* (linear relationship between *x* and *y*_1_). In Case II, we have *y*_2_= 5 (*x* − 0.5)^2^ + *ε* (nonlinear relationship between *x* and *y*_2_) and for Case III, *y*_3_ = 5 (*x* − 0.5)^2^ + 2*x* + *ε* (combination of linear and nonlinear relationships between *x* and *y*). Noise ε is a Gaussian distribution with a mean of zero.

We need to ensure the NMI estimation is robust because we are using windowed NMI, which involves a smaller number of time points. To study the impact of the number of data points in the NMI estimation, in each case, we took sample points x_i_ and y_i_ of size 35, 50, 75, 100 and measured their relationship before and after removing linear correlation. The dependency before removing linear correlation is represented as MI_1,_ and dependence after removing linear correlation is measured and represented by MI_2_ (Table 1).

**Table 1.** Assessing NMI performance on several samples with 35, 50, 75, and 100 elements in each set for linear, explicitly nonlinear, and combination dependencies. MI_1_ denotes the dependence before linear relationship removal, and MI_2_ represents dependency after linear removal. As the number of sample points gets smaller, the result goes further from ground truth, but the differences between before and after linear removal are still distinguishable.

|  | T=35 | T=50 | T=75 | T=100 | T=10,000<br>(ground truth) |
| --- | --- | --- | --- | --- | --- |
| **Case I: $y_1 = 2x + \epsilon$**<br>**(linear relationship)** | $MI_1 = 0.2399$<br>$MI_2 = 0.0088$ | $MI_1 = 0.2765$<br>$MI_2 = 0.0276$ | $MI_1 = 0.2875$<br>$MI_2 = 0.0206$ | $MI_1 = 0.3183$<br>$MI_2 = 0.0133$ | $MI_1 = 0.3305$<br>$MI_2 = 0.0127$ |
| **Case II: $y_2 = 5(x - 0.5)^2 + \epsilon$**<br>**(nonlinear relationship)** | $MI_1 = 0.1211$<br>$MI_2 = 0.1216$ | $MI_1 = 0.1633$<br>$MI_2 = 0.1591$ | $MI_1 = 0.1515$<br>$MI_2 = 0.1535$ | $MI_1 = 0.1754$<br>$MI_2 = 0.1758$ | $MI_1 = 0.2404$<br>$MI_2 = 0.2402$ |
| **Case III: $y_3 = 5(x - 0.5)^2 + 2x + \epsilon$**<br>**(linear and nonlinear relationships)** | $MI_1 = 0.1558$<br>$MI_2 = 0.1176$ | $MI_1 = 0.1748$<br>$MI_2 = 0.1385$ | $MI_1 = 0.2431$<br>$MI_2 = 0.1606$ | $MI_1 = 0.2419$<br>$MI_2 = 0.1777$ | $MI_1 = 0.3001$<br>$MI_2 = 0.2357$ |

### 2.3. Participants and Preprocessing

We used the fBIRN dataset analyzed previously used in (Damaraju et al.). The final curated dataset consisted of 163 healthy participants (mean age 36.9, 117 males; 46 females) and 151 age- and gender-matched patients with schizophrenia (mean age 37.8; 114 males, 37 females). Eyes-closed rsfMRI data were collected at seven sites across the United States (Keator et al., 2016). Informed consent was obtained from all subjects before scanning by the Internal Review Boards of affiliated institutions. Imaging data of one site was captured on a 3- Tesla General Electric Discovery MR750 scanner, and the rest of the six sites were collected on 3-Tesla Siemens Tim Trio scanners. Resting-state fMRI (rsfMRI) scans were acquired using a standard gradient-echo echo-planar imaging paradigm: FOV of 220 × 220 mm (64 × 64 matrices), TR = 2 s, TE = 30 ms, FA = 770, 162 volumes, 32 sequential ascending axial slices of 4 mm thickness and 1 mm skip.

Data preprocessed by using several toolboxes such as AFNI, SPM, GIFT. Rigid body motion correction was applied using the INRIAlign (Freire & Mangin, 2001) toolbox in SPM to correct head motion. To remove the outliers, the AFNI3s 3dDespike algorithm was performed. The rsfMRI data were resampled to 3 mm^3^ isotropic voxels. Then data were smoothed to 6 mm full width at half maximum (FWHM) using AFNI3s BlurToFWHM algorithm, and each voxel time course was variance normalized. Subjects with larger movements were excluded from the analysis to mitigate motion effects during the curation process.

### 2.4. ICA Analysis

The group ICA of fMRI toolbox (GIFT, http://trendscenter.org/software/gift) implementation of Group-level Spatial ICA was used to estimate intrinsic connectivity networks (ICNs). A subject-specific data reduction step was first used to reduce 162 time point data into 100 directions of maximal variability using principal component analysis. After PCA, the infomax approach (Bell & Sejnowski, 1995) was used to estimate 100 maximally independent components from the group PCA reduced matrix. To ensure the stability of the estimation, the ICA algorithm was repeated 20 times, and the most central run was selected as representative (Du, Ma, Fu, Calhoun, & Adali, 2014). Subject-specific spatial maps (SMs) and time courses (TCs) were obtained using the spatiotemporal regression back reconstruction approach (Calhoun, Adali, Pearlson, & Pekar, 2001; Erhardt et al., 2011) implemented in the GIFT software.

To label the components, regions of peak activation for each specific spatial map were obtained. After ICA processing, to acquire regions of peak activation, one sample t-test maps are taken for each SM across all subjects and then thresholded; also, mean power spectra of the corresponding TCs were computed. An independent component was identified as an intrinsic connectivity network (ICN) if its peak activation fell within gray matter and has low spatial overlap with known vascular, susceptibility, ventricular, and edge components corresponding to head motion. This results in 47 ICNs out of the 100 independent components.

The ICN time courses were detrended by removing linear, quadratic, and cubic trends and orthogonalized with respect to estimated subject motion parameters. Spikes were detected by AFNI3s 3dDespike algorithm and replaced by values of third-order spline fit. For more detail see (E. Allen et al., 2012; Damaraju et al., 2014). The fBIRN dataset obtained after processing resulted in a matrix of 159 time points × 47 ICNs × 314 subjects, including 163 Control and 151 SZ subjects. For more details, please see (Damaraju et al.).

### 2.5. Quantifying (Nonlinear) Dynamic Connectivity in fMRI Data

We compared HCs and SZs’ states of dynamic functional connectivity (dFNC), using both linear (Pearson correlation) and nonlinear (NMI approach as described in Section 0) dependencies. There are 47 ICN time courses of length 159 time points for each subject. From each time course x, a set of sliding windows x_t_, each of length 50 time points, is derived, that is 110 windows in total. To obtain linear dFNC, Pearson correlation between x_t_ and y_t_ is evaluated and resulted to 110 symmetric windowed-FNC matrices per individual. To quantify EN dFNC, for each t, the nonlinear dependency of pairs (x_t_, y_t_) are evaluated as described in Section 2.1. Quantifying Explicitly Nonlinear Dependency via a Normalized Mutual Information Approa, the linear dependency between x and y removed and then residual dependency is calculated by NMI. This procedure also resulted in 110 symmetric windowed-FNC matrices for each subject (Figure 2).

**Figure 2.**
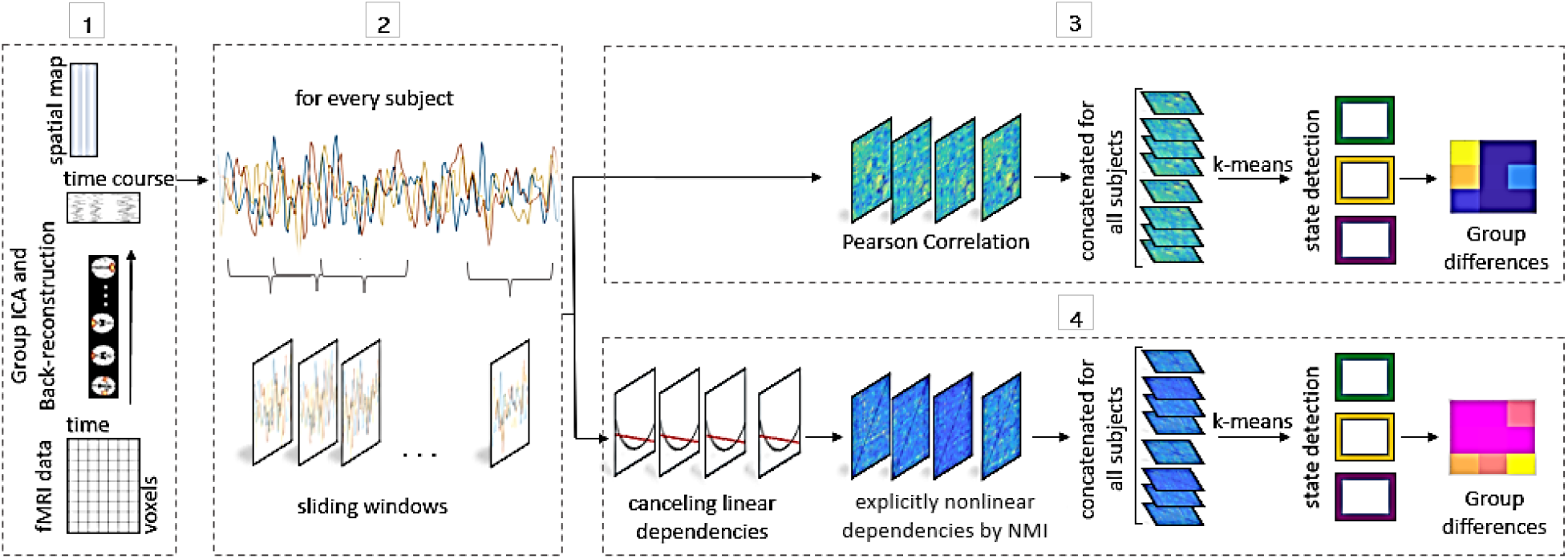
An overview of the linear and nonlinear dynamic FNC. 1) Group independent component analysis (ICA) is used to decompose resting-state data from 314 subjects into 100 components, 49 of which are identified as intrinsic connectivity networks (ICNs). Subject-specific spatial maps (SMs) and time courses (TCs) are estimated by the spatiotemporal regression back reconstruction method. 2) Sliding windows of length 50 time points are taken for each subject. 3) (Linear) Dynamic FNC is analyzed. First, the correlation matrices from windowed portions of each subject’s component TCs are assessed. Then the matrices are aggregated across all subjects and clustered by k-means clustering. Lastly, to probe the group differences between HC and SZ states are performed. 4) Nonlinear dynamic FNC is analyzed. Steps are identical as (3) except that linear correlation is removed for each window, and the remaining explicitly nonlinear dependencies are assessed by NMI. Matrices obtained from NMI are concatenated and grouped to states by using k-means clustering. Subject-specific state-types for each window are used to evaluate group differences.

We applied the k-means clustering method (using correlation distance) to obtain states for linear and nonlinear dFNC for cluster sizes of k = 2-10. The optimal number of distinct 3 dFNC states was estimated by conducting the elbow method. Final states are achieved by 100 repetitions, as shown in Figure 3.

**Figure 3.**
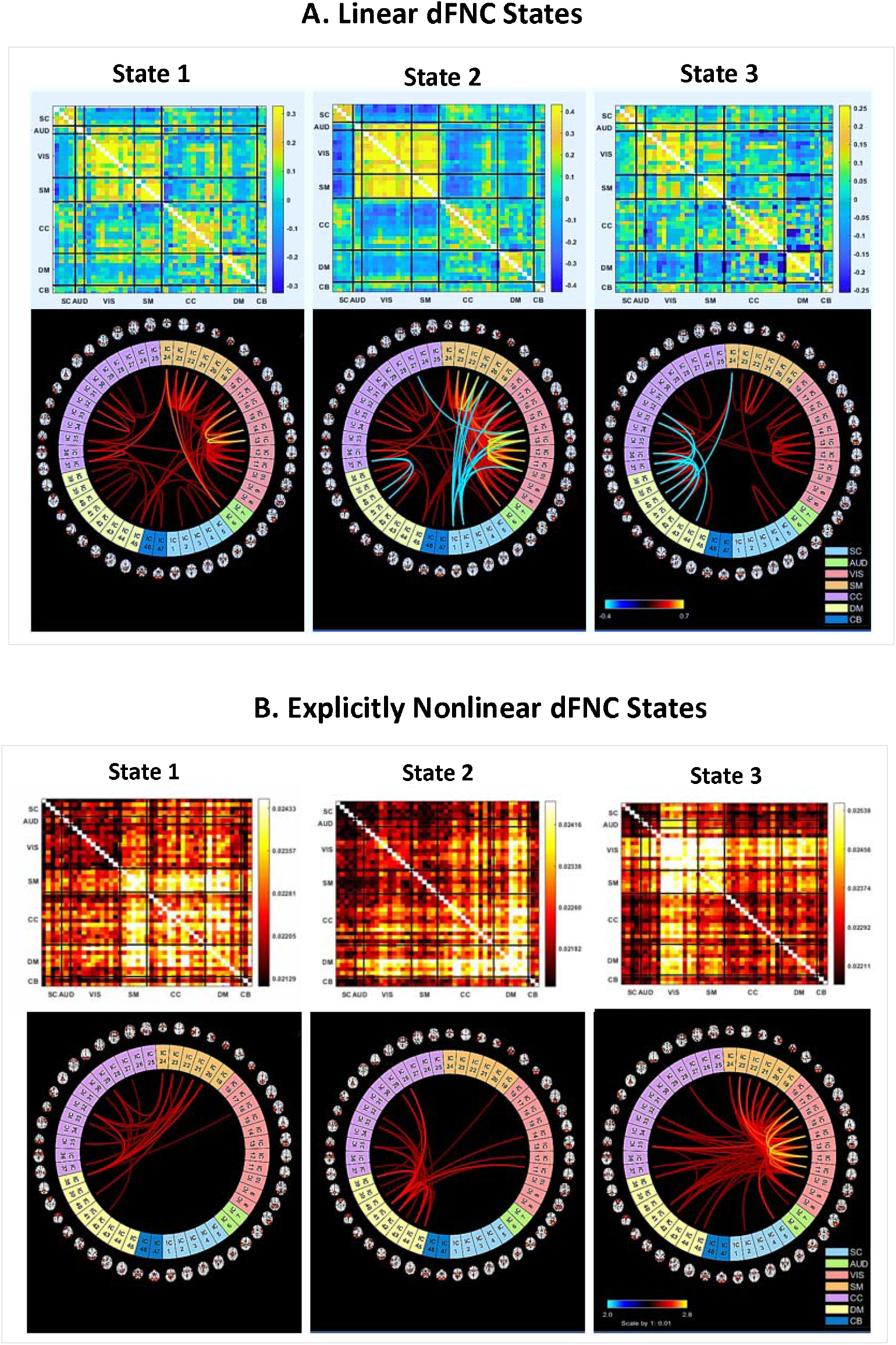
Linear and explicitly nonlinear states are acquired from the K-means clustering approach for k = 3. **A)** States achieved from clustering windows’ Pearson correlation of ICNs. The connectograms are thresholded at 0.3. **B)** States derived from clustering windows’ nonlinear dependencies of ICN’s. The connectograms are scaled by 1: 0.01 and thresholded at 0.025. Linear and nonlinear states demonstrate a distinctive contribution in and between networks. The rows of dFNC matrices were partitioned into sub-cortical (SC), auditory (AUD), visual (VIS), sensori-motor (SM), a broad set of regions involved in cognitive control (CC) and attention, default-mode network (DMN) regions, and cerebellar (CB) components.

After obtaining states, we assessed the group differences. Several quantities are computed at the level of individual’s window set and their corresponding K-means indices:

1. **Fractional Occupancy (FO)**- the percentage of overall time spent in each state.
2. **Dwell Time (DT)**- average duration of time spent in each state.
3. **Probability of Transitions (PT)**- the probability of transition from one state to other states.

Next, we compute a two-sample t-test to compare the differences of these results between controls and schizophrenia patients.

### 2.6. Relation between linear and nonlinear states

In this section, we are interested in finding out if there is any relationship between the three linear and nonlinear states. Several approaches are conducted. The contingency table of simultaneously being in each pair of states is calculated first for all individuals and then for HC and SZ separately (**Error! Reference source not found**.). The chi-square test rejects the null hypothesis that the linear and nonlinear dFNC’s states are independent (likewise in HC and SZ).

To further evaluate the group differences between the linear and nonlinear state vectors, a contingency table is computed for each individual. Then for each pair of linear and nonlinear states (9 pairs), the differences between HC and SZ are compared by a two-sample t-test.

## Results

### 3.1. Simulated Experiment

We examined 35, 50, 75, and 100 sample points on simulated data to find an applicable window length that captures nonlinear dependency by implementing the NMI method (Table 1).

In this work, we select 50-time points (100 s) from the result on simulation data (Table 1). This selection is based on two factors. The outcome of using sliding window analysis is sensitive to the length of the sliding window; in many studies, this length is between the 30s to 60s (E. A. Allen et al., 2014; Damaraju et al., 2014; Leonardi & Van De Ville, 2015), and in some cases longer (Leonardi & Van De Ville, 2015; Vergara, Abrol, & Calhoun, 2019). The other factor to consider is how small this length can be chosen. A small window size may result in the absence of nonlinearity. The NMI performance on various sample points in Table 1 shows that by taking 50 sample points, NMI can successfully distinguish between the linear and nonlinear relationships.

### 3.2. Results from fMRI Data

We measured linear and EN dynamic functional connectivity network (dFNC) of 163 healthy controls and 151 schizophrenia patients. The implementation of sliding window analysis and k-means clustering resulted in three states for each linear and nonlinear dFNC. Figure 3 shows dFNC states and their connectograms for better visualization of the nature of each state.

T-tests were used to identify group differences in several quantities of each linear and nonlinear states between controls and schizophrenic patients. The average fraction occupancy (FO) across healthy controls and schizophrenic patients of each state is calculated and listed in Table 2. The average dwell time (DT) across healthy controls and schizophrenic patients of each state is calculated and reported in Table 3.

**Table 2.** Fraction Occupancy of HC and SZ in linear and nonlinear dFNC. All linear states and nonlinear state 1 and 3 show highly significant group differences.

|  | State 1 | State 2 | State 3 |
| --- | --- | --- | --- |
| **Average across HC** | %30 | %43 | %27 |
| **Average across ZS** | %14 | %21 | %65 |
| **p-value** | $6.5383 \times 10^{-5}$ | $2.5613 \times 10^{-7}$ | $1.7319 \times 10^{-14}$ |
| **Average across HC** | %22 | %30 | %49 |
| **Average across ZS** | %34 | %37 | %29 |
| **p-value** | $1.2635 \times 10^{-4}$ | 0.0425 | $7.3181 \times 10^{-8}$ |

**Table 3.** Statical analysis of dwell time for linear and nonlinear states in HC and SZ. SZ spends significantly longer in linear state 3.

|  | State 1 | State 2 | State 3 |
| --- | --- | --- | --- |
| **Average across HC** | 54.69 | 59.08 | 68.35 |
| **Average across ZS** | 42.16 | 47.41 | 151.98 |
| **p-value** | 0.0177 | 0.0310 | $5.096 \times 10^{-12}$ |
| **Average across HC** | 13.15 | 12.98 | 19.52 |
| **Average across ZS** | 17.72 | 18.23 | 15.60 |
| **p-value** | 0.0147 | 0.0051 | 0.03244 |

For each subject, the probability of transition (PT) from state i to other states j, where j=1, 2, or 3, are calculated, i.e., the conditional probability p(“next state is j”|”now is in state i”). The comparison between healthy controls and schizophrenia patients is stated in (i, j) entry of the matrix in ..

Linear states show a high level of positive correlation within networks, and correlation between other networks fluctuates between negative and positive in each linear state. The graphical representations of linear dFNC states show dense interconnectivity within clusters but sparse (or negative in directed graphs) connections between nodes in different clusters. State 1 shows a considerable level of correlation within SC, AUD, VIS, SM, CC, DM networks, and between AUD, VIS, SM networks. However, State 2 shows a uniformly negative correlation between (AUD, VIS, SM) sets of networks and (CC) but a high level of positive correlation within networks and between AUD, VIS, and SM. State 3 shows a noticeably smaller range of correlation among all networks.

In the nonlinear dFNC states, we observe a high level of explicitly nonlinear dependencies within a specific network that contributes with broadly all other ICNs. Analogous graphical representations of states 2 and 3 are close to a star graph where only the center node is connected to other nodes. State 1 shows high EN dependency in SM and CC (based on the connectogram). State 2 shows the EN dependency between DM and other networks, and State 3 signifies substantial EN dependencies within VIS and SM and between other networks.

As it is shown in Table 2 and Table 3, HCs spend more time in the VIS and SM system (linear state 2 and nonlinear state 3) compared to patients that spend more frequently and longer in a low range of correlation (linear state 3) in SM, CC and DM networks (nonlinear state 1 and 2) where VIS network’s EN dependency has almost vanished.

With this view in Figure 6, HCs tend to spend longer and transit with a higher probability to EN state 3, which has higher explicitly nonlinear dependencies in VIS and SM. SZ spends longer in higher EN dependency of DM (state 2) but also transitions more to EN state 1 than 2. The average fraction occupancy for SZ in three explicitly nonlinear states is close to equal, while HC are comparably distributed uniformly. Unlike EN states, the average fraction occupancy of SZ is much higher in linear state 3.

## Discussion

Dynamic FNC provides a more natural way to analyze uncontrolled resting fMRI data and provide additional insight into brain activity (Hindriks et al., 2016; Hutchison, Womelsdorf, Allen, et al., 2013). However, virtually all research in this area, at least at the whole brain/connectome level, has focused on the linear correlation among time courses (E. Allen et al., 2012; Damaraju et al., 2014; Hutchison, Womelsdorf, Gati, et al., 2013; Obata et al., 2004). However, there is considerable evidence of nonlinearity in fMRI data (de Zwart et al., 2009; Sheth et al., 2004; Wan et al., 2006). Given that, in this work, we focus on the dynamics of explicitly nonlinear dependency among brain regions.

We measured explicitly nonlinear dFNC and dFNC between TCs obtained from processed fMRI data collected from HC and SZ in this work. Using a k-means classifier, each set of concatenated windows of either Pearson correlation or explicitly nonlinear dependencies are grouped into three integrated patterns of linear dFNC and explicitly nonlinear dFNC (Figure 3). These states are analyzed, and the differences between HC and SZ in each linear and EN state are compared and studied.

Our results suggest that EN states and linear states complement each other as their behavior has basic differences. For example, the average EN dwell time is considerably shorter than the linear dwell time (Figure 6). Also, the transition probability matrices are unique and reveal different information regarding HC and SZ (Figure 5). These unique aspects can also be observed in the average fraction occupancy.

In the linear states, we observe a strong positive correlation within networks such as (SC), (AUD, VIS, and SM), (CC), (DM), and (CB). For the relation between these networks, it is noted that the correlation swings between negative and positive in each linear state (Figure 3). Linear state 2 shows a sharp, clear, and intense pattern in terms of correlation between the networks. This pattern fades as moving to linear state 1 and becomes lowest in linear state 3.

For the explicitly nonlinear dFNC states, we measured NMI after removing linear correlation at each window. NMI takes values between 0 to 1. So, we lose the positivity and negativity interpretation here. Multi-network connections were observed in the EN compared to linear dFNC, one predominantly SM (with less but considerable level within CC and DM), one predominantly DM, one predominantly VIS and SM.

Our results (Figure 4, Figure 5, Figure 6) indicate that HC tend to have a high level of linear and EN dependencies (linear state 1, 2, and nonlinear state 3). At the same time, SZ tend to stay not only in a lower level of dependencies overall brain networks (linear state 3) but also in the absence of EN dependency in the VIS network (nonlinear state 1 and 2). However, SZ tend to have higher EN dependencies in CC and DM than HC.

**Figure 4.**
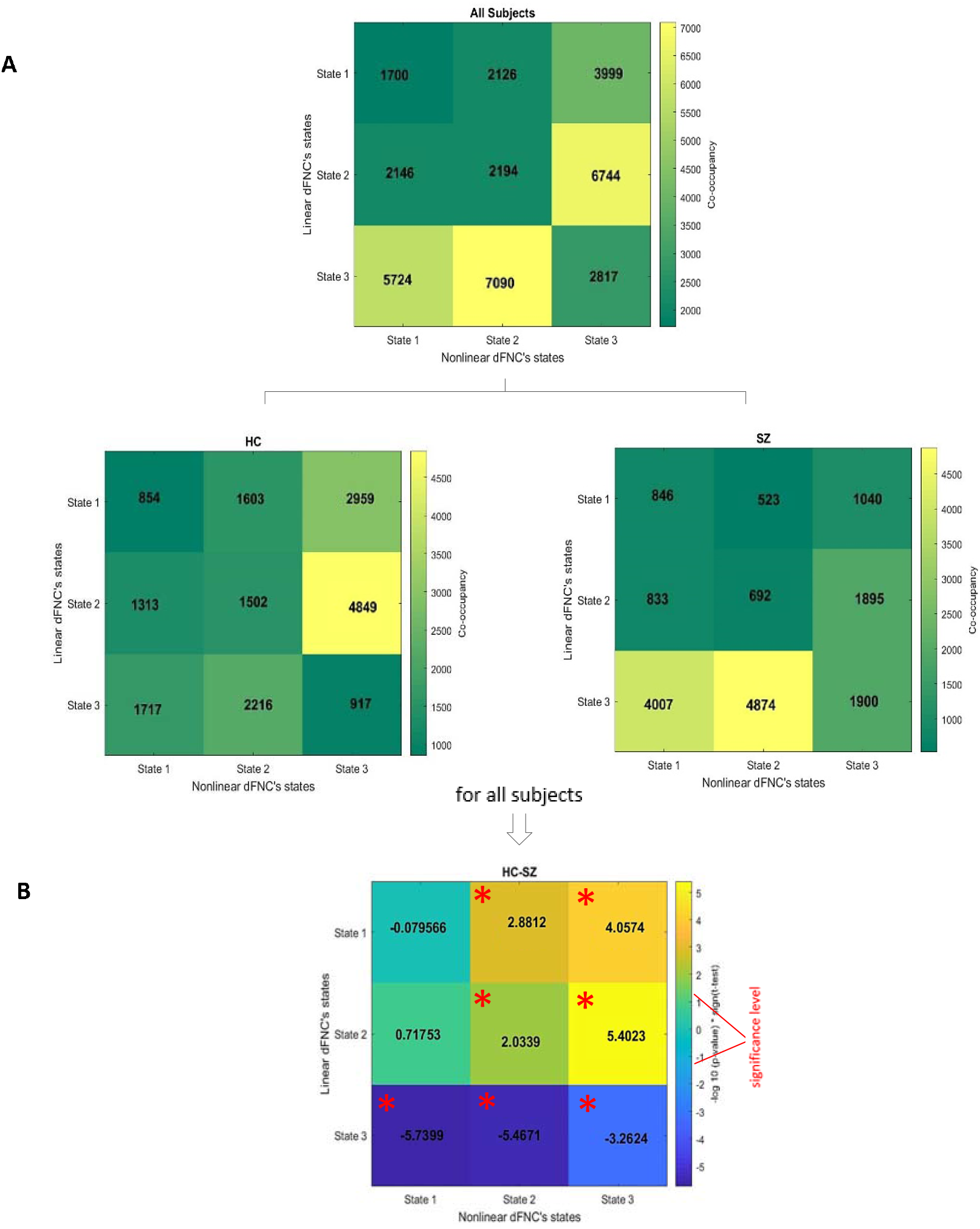
**A)** Contingency table of linear and nonlinear dFNC’s states for each group. **B)** FDR-adjusted p-values from comparing HC-SZ. Significant level indicated by red pointer for alpha= 0.05. HCs are observed significantly more in pairs (2,3) and (1,3), while SZs tend to be in (3,1) and (3,2) of linear-nonlinear states.

**Figure 5.**
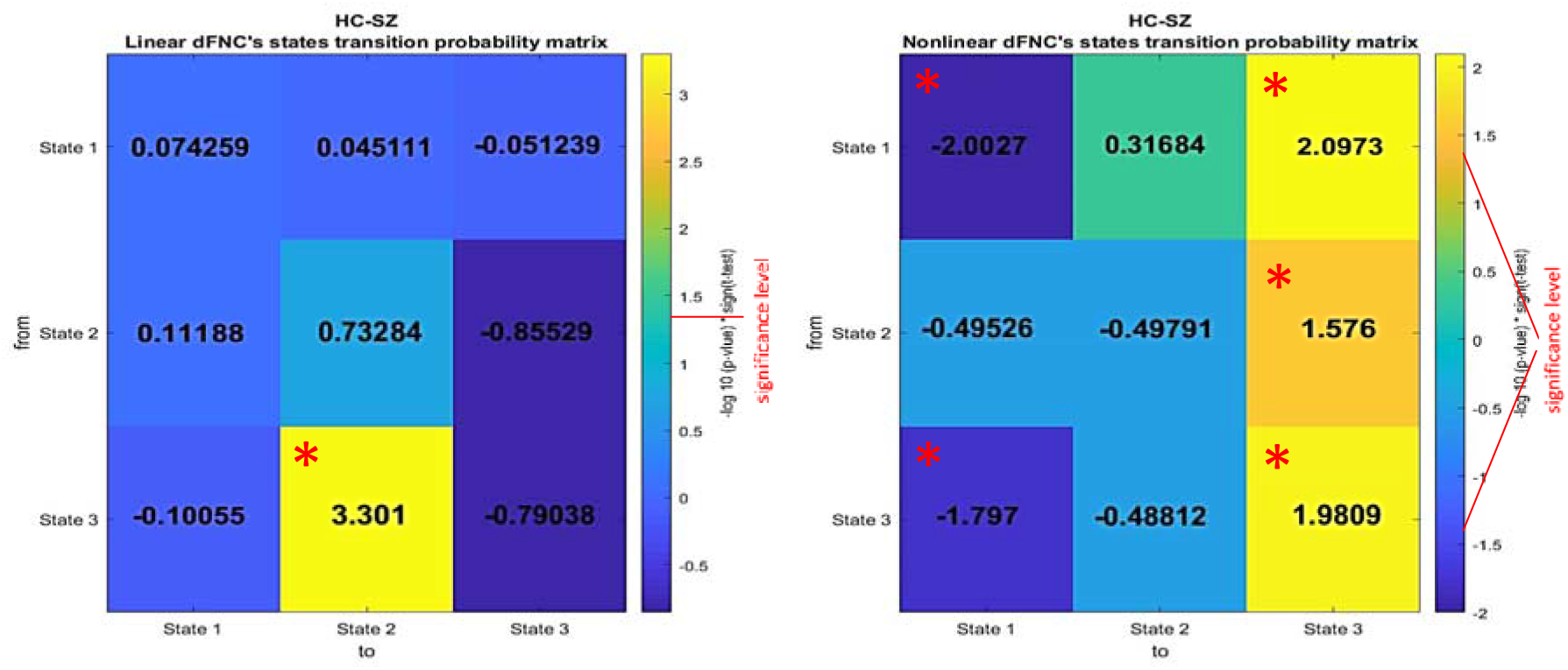
Comparing linear and nonlinear dFNC’s state transition. The probability of moving from state i to state j is calculated for each individual, and then for each pair (i,j), the difference between HC and SZ is measured and denoted in the (i,j) entry of the matrix. P-values are adjusted by FDR, and significant level indicated by red pointer for alpha= 0.05, and cells with significant value are denoted by asterisk. HC significantly tends to move to linear state 2 when they are in linear state 3. Also, they tend to move to nonlinear state 3, while SZ be inclined to move to nonlinear state 1. However, SZ tends to move and stay in linear state 3. They also show significantly transitions between nonlinear state 1 and also to nonlinear state 2.

**Figure 6.**
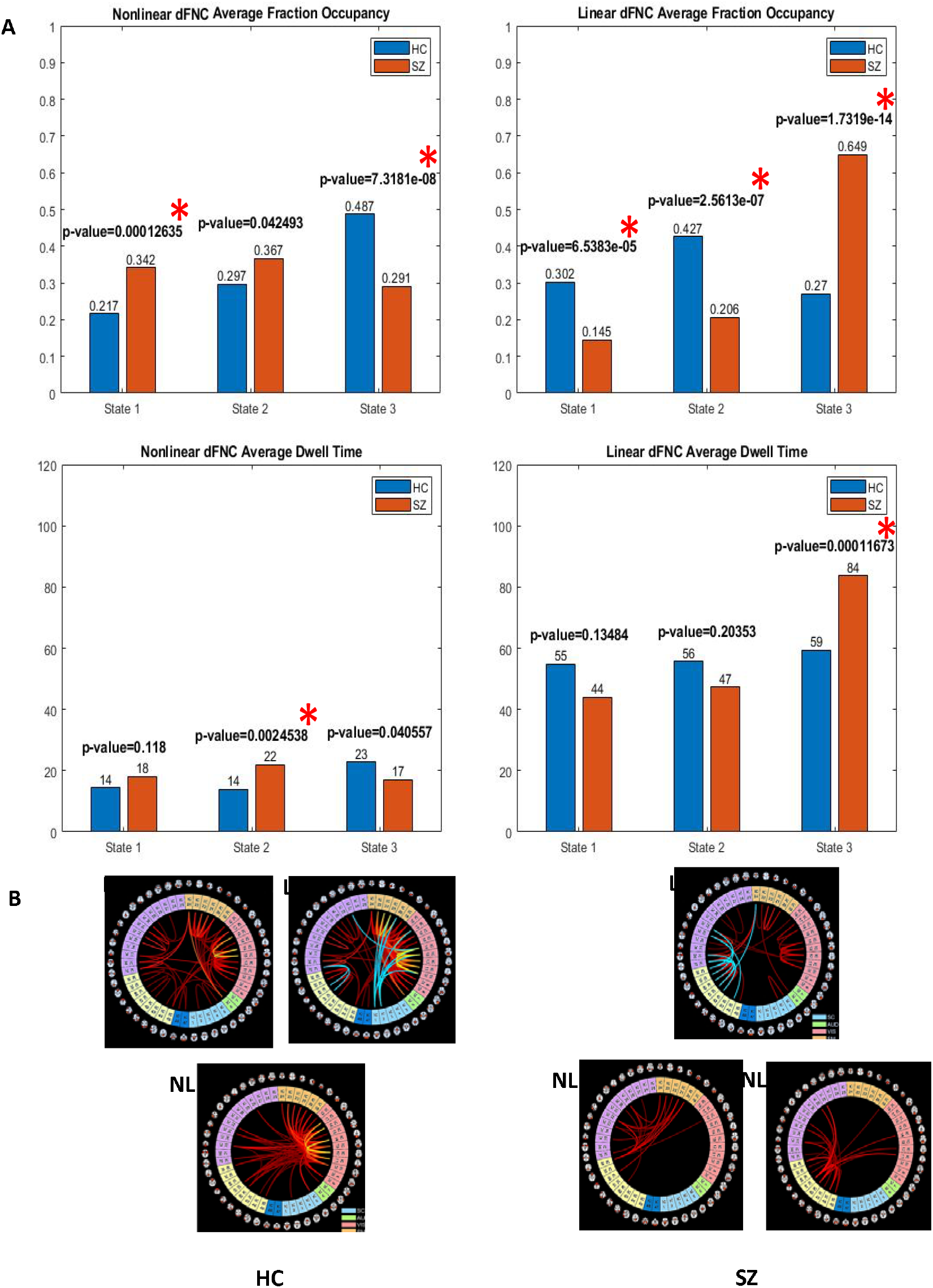
**A)** Another representation of fraction occupancy and dwell time. Asterisks denote significant p-values. **B)** Linear and nonlinear states that HC (left) and SZ (right) spend significantly more time in.

These results agree with our earlier findings in evaluating (static) EN FNC (Motlaghian et al., 2021), which indicated significant differences in EN dependencies within and between AUD, VIS, CC, and DM networks in HC and SZ over the entire run. Studying explicitly nonlinear dependencies in dynamic FNC helps unpack the informative structure of how temporally VIS and AUD networks are more active and CC is less involved in HC.

We also analyzed the relationship between fraction occupancy, dwell time of EN states, and symptoms of SZs, and didn’t find a significant relationship.

Measuring explicitly nonlinear dependencies by NMI is not symmetric, respecting the order of TCs. However, the preference is for being symmetric because we expect to observe the same amount of dependency regardless of the order of inputs. We addressed the symmetricity by taking the average between two results in our work. Another limitation of using NMI is that the interpretation of positivity and negativity is missed compared to the correlation.

For future work, more investigation of the impact of the window size on the sliding window analysis may drive more information about the dynamic states. The simulation demonstrated NMI showed good performance before and after removing the linear correlation with as few as 35 sample points. In our work, we used 50 time points which can be reduced without losing the NMI sensibility. It would also be interesting to utilize a filter bank approach to cover a larger range of window sizes/frequencies (Faghiri, Iraji, Damaraju, Turner, & Calhoun, 2021).

## Appendix A

### Supplementary Data

Supplementary data to this article can be found online at http://dx.doi.org/10.1016/j.nicl.2014.07.003

